# Targeted Protein Degradation via Nanoparticles

**DOI:** 10.1101/2022.09.21.508905

**Authors:** Yang Liu, Runhan Liu, Jiawei Dong, Xue Xia, Haoying Yang, Sijun Wei, Linlin Fan, Mengke Fang, Yan Zou, Meng Zheng, Kam W. Leong, Bingyang Shi

## Abstract

Strategies that hijack selective proteins of interest (POIs) to the intracellular protein recycling machinery for targeted protein degradation (TPD) have recently emerged as powerful tools for undruggable targets in biomedical research and the pharmaceutical industry. However, targeting any new POI with current TPD tools requires laborious case-by-case design for different diseases and cell types, especially for those extracellular targets. Here, we observed that nanoparticles (NPs) can mediate the receptor-free internalization of hijacked protein and further developed a generic paradigm for both intra- and extracellular POI degradation, by making full use of clinically approved components. The phenomenon is general, as we found nanostructures such as lipid nanoparticle (LNP), liposomes, exosomes, polymeric nanoparticles, inorganic nanoparticles and their hybrid nanoparticles modified with POI-recognizing moiety (antibody, peptides, small molecule drugs) can mediate TPD for a wide range of extracellular/membrane and intracellular targets. The super flexible and feasible-to-synthesize TPD-NPs paradigm may revolutionize the current TPD tools development landscape and it provides fundamental knowledge to receptor mediated drug therapies.

**Highlights:**

- Nanoparticle mediated targeted protein degradation (TPD-NP) can be constructed by “Mix-and-Match” and do not require *de novo* synthesis or specific internalization design.
- TPD-NPs can be equipped with specific cell type targeting capacity, loading and controlled release of therapeutic cargos, as well as biological barrier penetration capacity.
- Assembling components can be clinical-approved or biodegradable for translational medicine.
- TPD-NP highly boosted current application platforms of nano-delivery and TPD.

## INTRODUCTION

Targeted protein degradation (TPD) selectively degrades proteins of interest (POIs) by the intracellular protein recycling machinery through the ubiquitin–proteasome system (UPS) and endolysosomal or macroautophagic pathways ^1^. TPD holds great promise for currently undruggable protein targets that traditional pharmaceutical approaches fail to satisfactorily address. ^2^ While current TPD platforms have attracted profound interest over the past two decades, they each have unresolved limitations, and a new technological paradigm shift is required to fulfill the promise of TPD. First-generation TPD is UPS dependent ^3^, PROTAC is the best-developed first-generation TPD and has been evaluating in clinical trials ^4^. Second-generation TPD tools appeared quite recently which exploit the intracellular autophagy machinery and mediate TPD independently of the UPS mechanism ^5-7^. All these pioneering studies highlighted potentials to target ‘undruggable’ proteins. However, most of these TPD platforms were designed for cytosol-localized protein targets. To achieve targeting of noncytosolic targets for degradation, lysosome-targeting chimeras (LYTACs) ^8^ were developed to bind and direct membrane-associated to lysosomes for TPD. LYTAC formulas are composed of an antibody or small molecule tagged with a mannose-6-phosphonate-rich tail, which is a CI-M6PR-binding ligand. CI-M6PR can therefore drive the antibody- or small molecule-hijacked POI complex to the lysosome and initiate the degradation of disease-relevant extracellular or membrane proteins ^8^. Extracellular TPD tools are still poorly investigated, existing examples ^9-12^ with similar designing rationale with LYTAC concluded a successful TPD tool aiming to degrade extracellular proteins requires the following essential components: i) a specific binder for POI hijacking; ii) a cell-penetrating ligand for POI internalization, and iii) a motif that can navigate the POI complex to the protein recycling center for POI degradation. However, applying the abovementioned TPD formulations to target any new POI requires laborious case-by-case design for different diseases and cell types. Therefore, a generic and simple design strategy of TPD tools exempting the abovementioned prudish design requirements is desirable.

Biocompatible NPs have been used for drug delivery in clinical practice for decades including the most recent COVID-19 mRNA nanovaccines. Their structural flexibility and facile surface potentiality are attractive properties for developing versatile paradigm as a TPD tool ^13^. Most importantly, engineered NPs can be taken up through multiple endocytosis processes after administration ^14^, entering the cell without cell-type specific receptors facilitating transport. After endocytosis, NPs are trafficked to acidic environments that mature into endosomes, which are then directed to lysosomes ^15^. No need for specific receptors to facilitate cell entry (i.e., receptor-free entry) and lysosome-directed capability, combined with the surface decoration potential of NPs. All these indigenous superiorities suggested that engineered NPs have the potential to bypass all the obstacles faced by current TDP formulations. In this study, we present a generic strategy of TPD-NPs for designing and developing effective TPD tools for both intra- and extracellular POI degradation as a paradigm which may revolutionize the current TPD tools development landscape and broadens the application of nanomedicine in an unexpected direction due to the superiorities in structural flexibility, synthetic practically and working potency.

## RESULTS

### EGFR degradation through antibody-linked PLGA TPD-NPs

We chose PEG-DSPE, polyethyleneglycol-(1,2-distearoyl-sn-glycero-3-phosphoethanolamine-N-[methoxy(polyethyleneglycol)]), as a POI binding linker since PEG-DSPE is highly biocompatible and has been approved by the U.S. Food and Drug Administration (FDA) for drug delivery ^16^. Antibodies are chosen as the POI binder, considering their high affinity to POIs and the fact that antibody‒drug constructs are well accepted clinically ^17^. PLGA, poly(lactic-co-glycolic acid) copolymer, the most extensively studied and FDA-approved therapeutic vehicle, was used for assembly with the antibody-PEG-DSPE arm (Figure 1A). We then decorated the TPD-NP with therapeutic epidermal growth factor receptor (EGFR) antibody monoantibody nimotuzumab (NTZ) to form first NPs for TPD application (TPD-NP) as proof-of concept (Figure 1A), considering that EGFR is a crucial tumor progression signaling transducer ^18^ on the cell surface, and assessed whether tumor cell EGFR could be degraded. The NTZ-NPs were approximately 150 nm in diameter (Figure 1B), and the successful linking between the antibody and polymer was visualized by native gel (Figure 1C). EGFR is highly expressed by MDA-MB-231 human breast cancer cells and provides a suitable model for assessing the effects of NTZ-NP on EGFR protein degradation. Treatment with NTZ-NP induced EGFR TPD (Figure 1, D and E). A similar TPD effect was observed when NTZ was replaced by cetuximab (CTX), a classic human-mouse chimeric EGFR monoantibody, in the NP (Figure S1A). We also applied a GFP tag fused to the intracellular domain (C-terminal before stop codon) of EGFR to facilitate tracing of EGFR internalization and degradation. Time-lapse imaging showed rapid EGFR removal from the plasma membrane within 24 h after treatment (Figure 1F), which peaked between 24 h and 48 h post-treatment (Figure 1F, Figure S1, B and C; Movie S1). When treatment was interrupted after 24 h, by refreshing the culture medium, expression of the POI gradually recovered (Figure S1B), in line with the temporary degradation profile of the TPD mechanism. Indeed, after reincubation with fresh medium, no obvious protein dysregulation was detected by protein native gel assay (Figure S1D). Surface EGFR degradation was also verified by confocal imaging (Figure 1G). Degradation of POI is dose dependent, and the TPD-NP dose can reach 1 nM (Figure S1E), which is competitive with the 10 nM minimal dose of the LYTAC tools ^8^. Furthermore, to demonstrate functional EGFR protein degradation, the expression of proteins downstream of EGFR was evaluated. Upregulation of active phosphorylated-AKT (p-AKT) as a complementary HER3/IGF1 downstream pathway has been reported in response to EGFR degradation ^19^ and was observed after NTZ-NP treatments (Figure S1F). MDA-MB-231 tumor cell proliferation inhibition also consequently occurred (Figure 1H). Taken together, these data suggested that TPD-NP is sufficient to degrade targeted surface proteins through an internalization-and-degradation process (Figure 1I).

**Figure 1.**
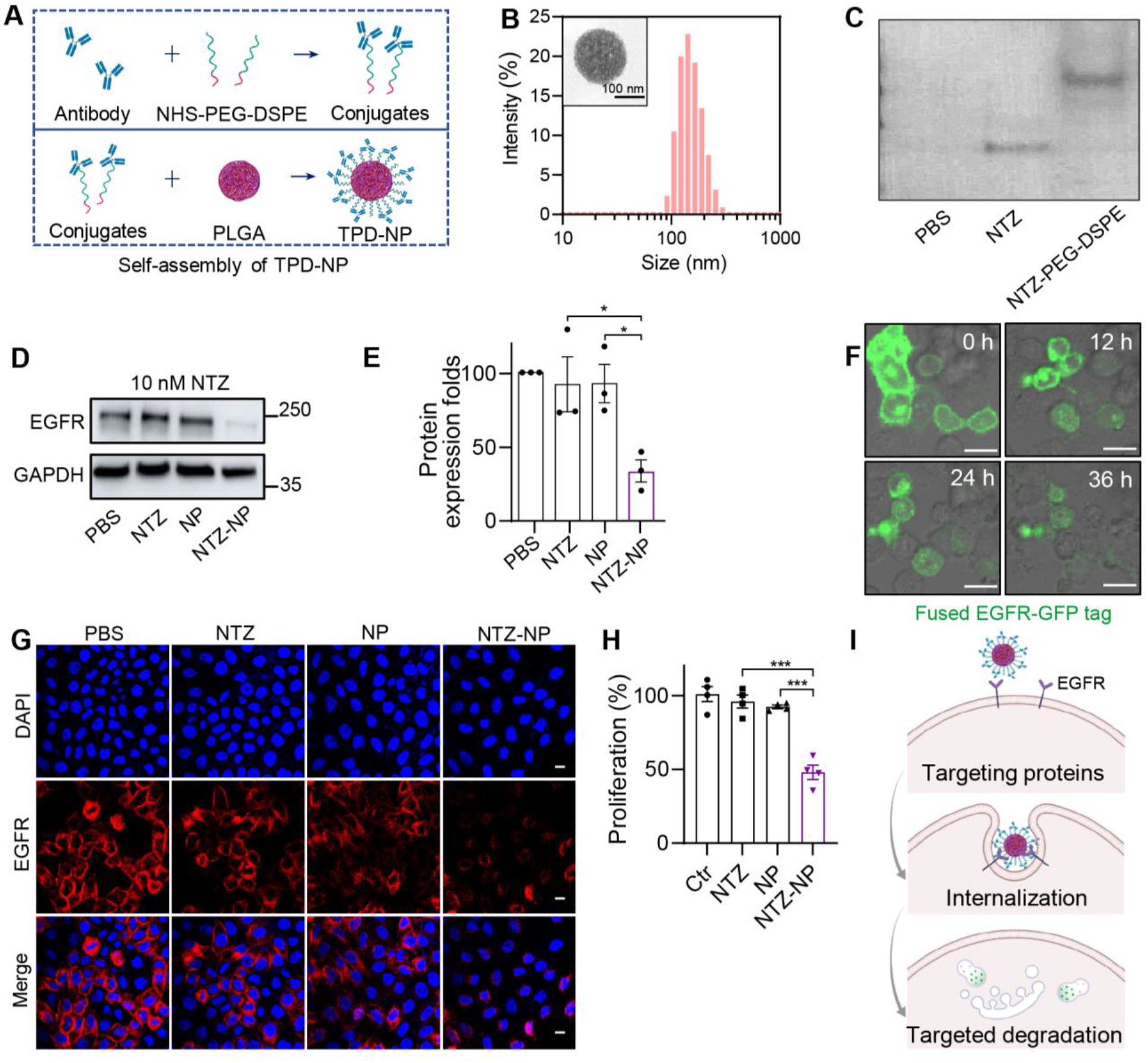
EGFR degradation by antibody-linked PLGA nanoparticles. (**A**) Schematic representation of the antibody-based TPD-NP structure by self-assembly of the hybrid NP. The PLGA core was coated with a PEG-DSPE-linked antibody. (**B**) Size distribution and TEM micrograph of the TPD-NPs. NTZ was used for surface decoration. Scale bar = 100 nm. (**C**) Native gel with Coomassie staining of the free antibody and PEG-DSPE linked antibody. (**D and E**) EGFR protein expression in MDA-MB-231 cells treated with NTZ-NPs for 24 h and statistics from 3 biological replicates. TPD-NP group contained an equivalent antibody dose of 10 nM. (**F**) Degradation of cell surface EGFR in MDA-MB-231 cells as determined in live cells after treatment with 500 nM equivalent antibody dose of NTZ-NP. Scale bar = 25 μm. (**G**) Confocal imaging of EGFR degradation in MDA-MB-231 cells after treatment with a 500 nM equivalent antibody dose of NTZ-NP for 24 h. Scale bar = 10 μm. (**H**) Inhibition of cell proliferation in HepG2 cells after treatment with NTZ-NP (500 nM, 24 h). n=4, mean ± s.e.m. *t* tests. \**P*<0.05, \*\*\**P*<0.001. (**I**) Schematic illustration of EGFR degradation using TPD-NPs.

### Broad-spectrum application of TPD-NP across different diseases, POI motifs and NP formulations

As a cross-check, we further assessed EGFR degradation in other tumor cell types (HeLa cervical cancer cells and U87 glioma cells). EGFR degradation was observed after treatment with NTZ-NP in these cell lines (Figure S2A). We then aimed to verify and expand the applications of the TPD-NP strategy to more clinically relevant targets (Figure 2A). As tumor immunotherapy is one of the most promising new cancer therapies, we assessed whether TPD-NP strategies could induce inhibition of the PD-1 and PD-L1 pair that facilitate tumor immune escape ^20^. Therefore, we applied clinically used immune checkpoint-inhibiting monoantibodies targeting PD-1 and PD-L1, camrelizumab (CRLZ) and atezolizumab (ATZ), respectively, to our TPD-NP strategy. The data showed that CRLZ-NPs can degrade PD-1 in PD-1-expressing Jurkat cells, while ATZ-NPs induced a decrease in PD-L1-expressing 293T cells (Figure 2, B and C, Figure S2B). Additionally, we assessed the potential of TPD-NP to knock down HER2, an established therapeutic target in cancer expressed in a large subset of patients ^21^. The clinically used therapeutic monoantibody pertuzumab (PTZ) targets HER2 by binding to the dimerization domain of HER2, thereby inhibiting HER2 heterodimerization ^21^. PTZ-linked NPs (PTZ-NPs) also successfully degraded HER2 expression in 293T or HepG2 cells (Figure 2, D and E). Considering that bispecific antibodies have emerged as an important new pursuit in the pharmaceutical industry, we also designed and tested INE (HER2 therapeutic antibody inetetamab)-NTZ-NP as a dual-targeting TPD-NP. PTZ/NTZ-NPs achieved effective protein degradation of both EGFR and HER2 (Figure 2F), confirming that our facile approach allows the construction of bespoke dual-targeting NP solutions by simple mix-and-match.

**Figure 2.**
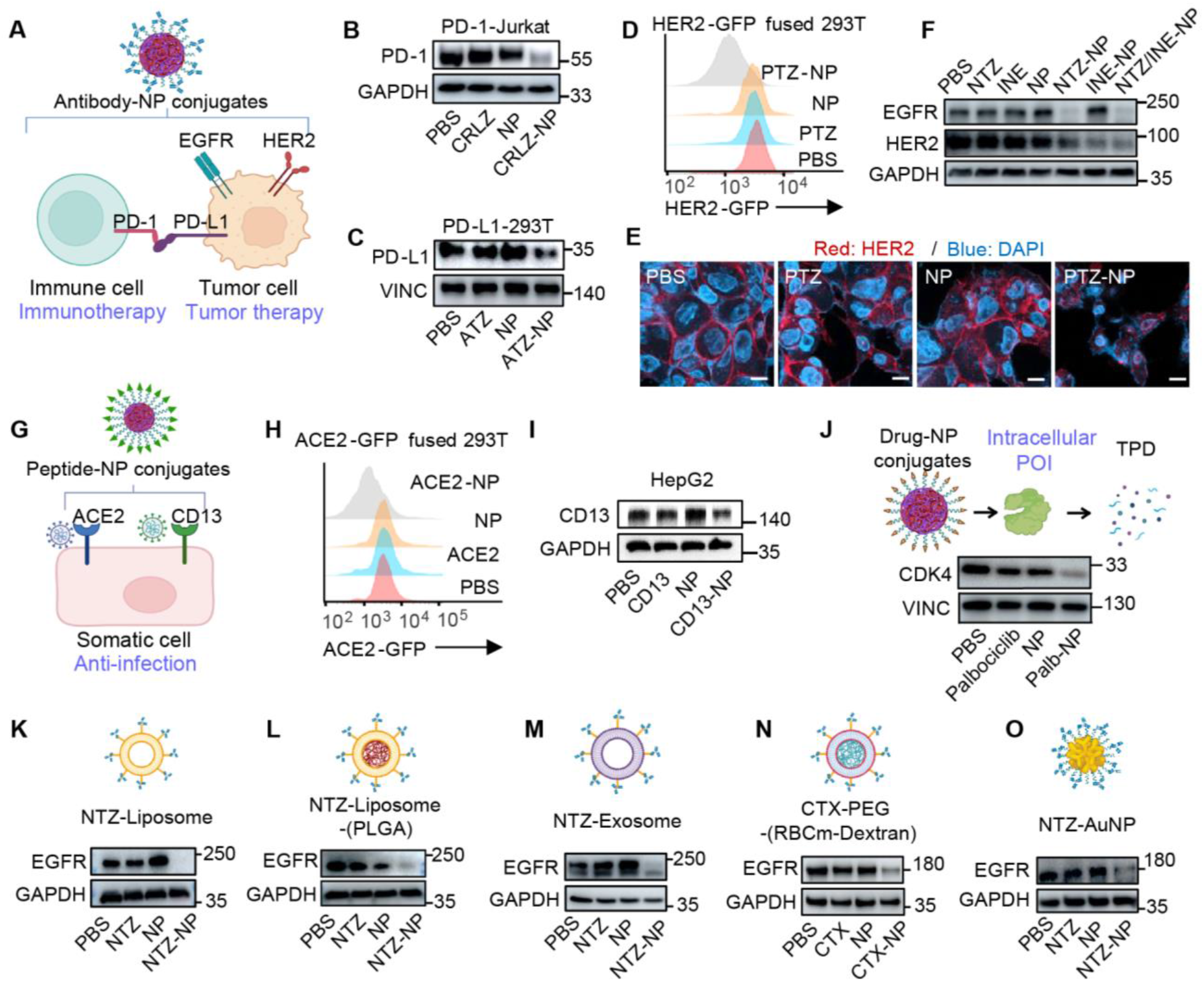
Broad-spectrum application of TPD-NP across different diseases, POI motifs and NP formulations. (**A**) Schematic summary of the representative POI and antibody-NP conjugates formulated for TPD evaluation. (**B**) PD-1 expression levels in Jurkat cells overexpressing PD-1 after treatment with 500 nM CRLZ-NP for 24 h. (**C**) PD-L1 expression levels in 293T cells overexpressing PD-L1 after treatment with 500 nM ATZ-NP for 24 h. (**D**) Flow cytometry data for HER2-GFP fusion protein expression levels in 293T cells after treatment with 500 nM PTZ-NP for 24 h. (**E**) Confocal imaging of HepG2 cells 24 h after PTZ-NP treatment. Scale bar=10 μm. (**F**) HER2 and EGFR expression levels in 293T cells after treatment with 500 nM NTZ-NP, 500 nM INE-NP and NP conjugated with both NTZ and INE for 24 h. (**G**) Schematic illustrating that the POI targeting moiety was replaced by ACE2 and CD13 binding peptides. (**H**) ACE2-GFP fusion protein expression in 293T cells (GFP fused to intracellular domain, C-terminal), after treatments with 10 μM ACE2 binding peptide (ACE2) or ACE2-NP for 24 h. (**I**) CD13 expression levels in HepG2 cells after treatments with 2 μM CD13 binding peptide (CD13) or CD13-NP for 24 h. (**J**) Schematic illustrated the POI targeting moiety was replaced by small molecule and targeted POI changed to intracellular protein. CDK4 protein levels after treatment with 2.5 μM palbociclib-NPs for 24 h. (**K**) EGFR expression after 500 nM NTZ-liposome treatments for 24 h in MDA-MB-231 cells. (**L**) EGFR expression after 500 nM NTZ-(Liposome-PLGA) NP treatments in MDA-MB-231 cells for 24 h. (**M**) EGFR levels in MDA-MB-231 cells after treatment with 1 μM NTZ-Exosome for 24 h. (**N**) EGFR expression after 500 nM CTX-(red blood cell membrane-acetal-dextran) NP treatments in MDA-MB-231 cells for 24 h. (**O**) EGFR levels in MDA-MB-231 cells after treatment with 1 μM NTZ-AuNPs for 24 h. POI binding moieties in all TPD-NPs were linked to NPs by PEG.

We next determined whether replacing antibodies with peptides or small molecules can achieve equivalent TPD effects. We applied a specific binding peptide of angiotensin-converting enzyme 2 (ACE2), one of the core surface receptors of SARS-CoV-2 ^22^. Inhibition of ACE2 has been suggested to be a broad-spectrum coronavirus blocker ^23^. We also constructed a TPD-NP incorporating CD13 (aminopeptidase N), a tumor surface marker relevant to tumor proliferation that also acts as a common-cold hCoV-229E coronavirus receptor ^24^ (Figure 2G). To monitor the degradation of the surface protein ACE2 by engineered peptide NPs, we employed GFP tags. Both ACE2 and CD13 were degraded by their corresponding TPD-NP systems (Figure 2, H and I, Figure S2B). We also wondered if TPD-NP could target intracellular POIs. To characterize this, the intracellular protein cyclin-dependent kinase 4 (CDK4)-binding chemical drug palbociclib (Palb), which was approved by the FDA for tumor therapy, was conjugated to PEG-DSPE to generate Palb-NPs. Treatment with Palb-NPs showed robust CDK4 degradation (Figure 2J). Collectively, these data suggest that the TPD-NPs may be applicable to a broad spectrum of therapeutic purposes, including tumor therapy, immune modulation, prevention of somatic cell infection and many other pathological conditions, with in-scope targeted POIs enlarged to encompass intracellular and extracellular proteins.

We next explored the range of applicable NP compositions that could suit TPD applications. In addition to PLGA, lipid based nanostructures including liposome, lipid nanoparticle (LNP) and lipopolyplex are the best-developed medically used NP formulations, having first been clinically applied in 1995 ^25^. Accordingly, we assembled NTZ-PEG-DSPE into the classic clinically used cholesterol liposome and lipopolyplex ^25^, respectively forming NTZ-Lipo (Figure 2K) and NTZ-lipopolyplex (Figure 2L). All these lipid-based NPs showed sufficient EGFR degradation at the tested doses (Figure 2, K and L). This inspired us that this phenomenon may be fundamental to the majority of nanoparticle formulations. To verify this proposition, we engaged exosomes, membrane camouflaging NP and gold NPs (Au-NPs), and observed that these formulations also efficiently degraded EGFR protein (Figure 2M, N and O). Collectively, these data suggest that NP cellular internalization is a suitable process for TPD. This fundamental “kiss and eat” phenomenon may broaden the application of nanomedicine in an unexpected direction and may explain some relative biological process.

### Cellular internalization of TPD-NPs requires no additional cell-specific penetration facilitators and completes TPD through autolysosomes

Next, we sought to determine the underlying mechanism(s) of the TPD-NP platform after we observed successful outcomes of TPD by a range of TPD-NP designs and in different target cell lines. First, understanding how the predegradation complex (POI-binder-NP) is internalized into cells is crucial for understanding why TPD-NP differs from other existing TPD platforms. An important point to emphasize is that for other recently developed extracellular protein degraders, a carefully designed cell-specific cell penetration facilitator is essential to enable the protein hijacking complex to enter the cell ^3^. However, a key significant advance of NP-based TPD tools is that no cell penetration facilitators are needed, and cellular uptake mechanisms reflect recently disclosed processors ^14^ in a process called enhanced permeability and retention (EPR). EPR describes the internalization of an NP via clathrin- or dynamin-dependent endocytosis, which forms an endosome or uptake by micropinocytosis ^26^ (Figure 3A). To identify the exact internalization pathway for POI-TPD-NP complexes, sucrose, methyl-β-cyclodextrin, nystatin and ethylisopropylamiloride (EIPA) were utilized to inhibit clathrin- and caveolae-dependent endocytosis or macropinocytosis, respectively ^26^. EGFR degradation in MDA-MB-231 cells was profoundly reversed by either sucrose or nystatin inhibitors (Figure 3, B and C, Figure S3, A to E) but not by EIPA treatment (Figure3D, Figure S3F). Successful inhibition of EGFR degradation confirmed that the POI-TPD-NP complexes could be taken up by cells via clathrin- and caveolae-dependent endocytosis.

**Figure 3.**
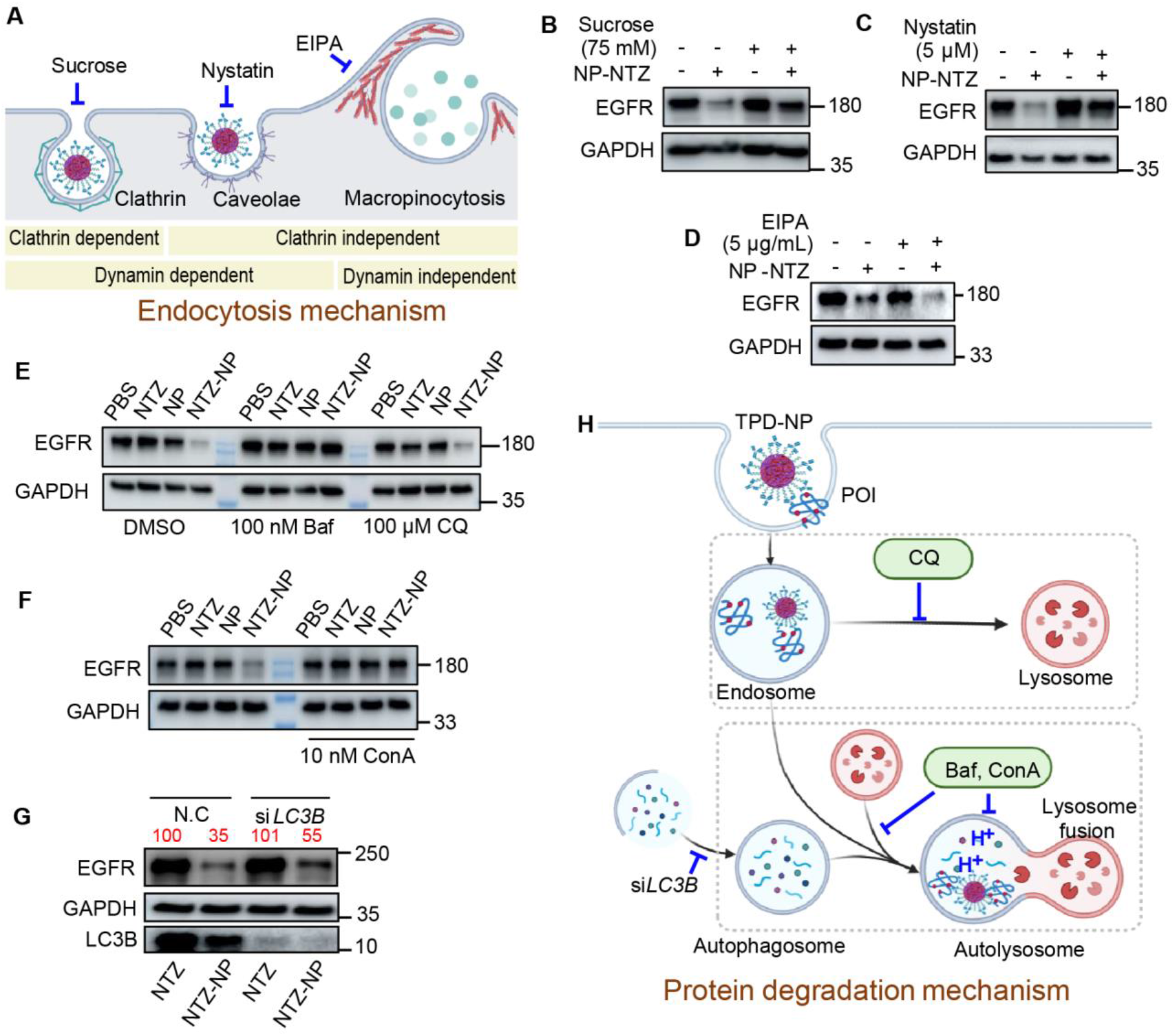
Engulfment of the POI-TPD-NP complex does not require specific cell penetration facilitators and achieves TPD through the autolysosome. (**A**) Schematic summary of the general cellular uptake pathways of NPs and representative inhibitors. (**B to D**) EGFR degradation in MDA-MB-231 cells after incubation with 500 nM NTZ-NP and 75 mM sucrose or 5 μM nystatin or 5 μg/mL EIPA for 24 h. (**E to F**) EGFR protein degradation evaluation in MDA-MB-231 cells after treatment with 500 nM NTZ-NP, together with the CQ, Baf and ConA for 24 h. (**G**) EGFR protein expression in MDA-MB-231 cells after treatment with 80 nM *LC3B* targeting siRNA (si*LC3B*) transfection and 500 nM NTZ-NP. (**H**) Schematic summary of the general non-cytosolic protein degradation pathways of TPD-NPs.

We then investigated the protein degradation mechanism, focusing on how the POI was automatically directed to the degradation center and which degradation pathway was engaged. We first tested whether lysosomes were involved in the degradation process by using the lysosomal inhibitors chloroquine (CQ) and bafilomycin A1 (Baf). These inhibitors were also used to reveal the LYTAC TPD mechanism ^8,10^. Our data indicated that Baf indeed significantly decreased TPD-NP-based EGFR degradation (Figure 3E), but to our surprise, CQ failed to reverse EGFR degradation over a wide dosage range (Figure 3E, Figure S4A). Previous investigations have shown that Baf can inhibit lysosome fusion to autolysosomes and serves as a specific and potent H^+^-V-ATPase inhibitor, thereby inhibiting the acidification of the autolysosome ^27^, indicating the possibility that TPD-NPs mediate POI degradation in autolysosomes. The experimental results with autolysosome acidification inhibitor, concanamycin A (ConA), further confirmed that the autolysosome is essential for the NP-mediated degradation process (Figure 3F, Figure S4B). To further confirm the autolysosome involvement, siRNA targeting *LC3B* was employed considering LC3B is one of the key player in autophagocytosis maturation ^28^, data suggested that si*LC3B* can indeed partially defect the TPD process (Figure 3G). While the ubiquitin–proteasome system (UPS) inhibitor MG132 showed no obvious impact on EGFR degradation in a wide concentration range (Figure S4C). Thus, we conclude that autolysosome is the key contributor to NP-mediated TPD (Figure 3H).

### TPD-NPs can efficiently degrade targeted proteins *in vivo*

One of the key highlights in the construction of our TPD-NPs was the use of clinically approved elements, suggesting that TPD-NPs have great potentials for rapid clinical translation. Given that the *in vivo* performance of previous extracellular TPD tools remained poorly investigated due to indigenous limitations, especially in solid tumors, we thereby assessed the *in vivo* performance of TPD-NPs by evaluating the antitumor effects of EGFR degradation. A subcutaneous tumor model was generated by transplanting MDA-MB-231 human breast cancer cells into nude mice. NTZ-NPs (15 mg/kg) were subcutaneously injected every second day for 7 cycles as depicted in the schedule (Figure 4A). TPD-NP treatment resulted in inhibition of *in vivo* tumor progression as determined by tumor volume (Figure 4B). Subsequently, the tumor was harvested, and EGFR protein expression and apoptosis-related downstream proteins were evaluated. These data demonstrated that NTZ-NP can sufficiently decrease EGFR expression *in situ* (Figure 4, C and D, Figure S5), providing the first direct evidence of protein degradation in solid tumors by an extracellular TPD platform. Additionally, as a consequence of EGFR inhibition, the expression of the tumor core apoptosis effector cleaved caspase3 was elevated (Figure 4C). Notably, treatment with TPD-NP induced a greater degree of tumor inhibition than antibody treatment alone (Figure 4B) with negligible body weight changes during treatments (Figure S6), indicating the therapeutic potential of TPD-NP-mediated protein degradation.

**Figure 4.**
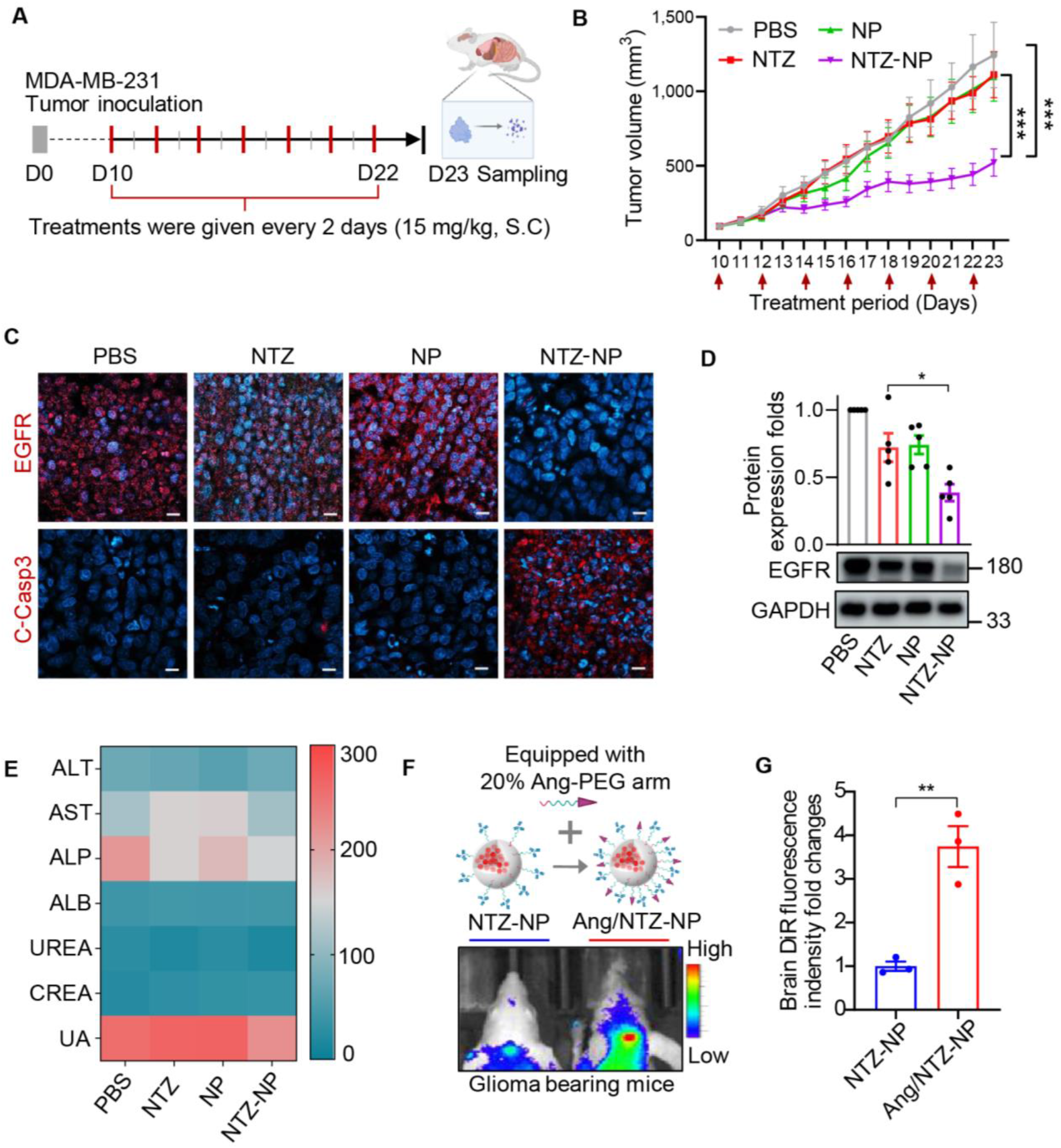
TPD-NPs can sufficiently degrade targeted proteins in solid tumors *in vivo*. (**A**) Treatment schedule of EGFR degradation evaluation in the subcutaneously transplanted MDA-MB-231 tumor animal model. The treatment protocol was 15 mg/kg NTZ-NPs in PBS injected subcutaneously for 7 cycles every second day. (**B**) Daily tumor volume measurements after treatment. n=10, mean ± s.e.m. (**C**) Immunostaining of EGFR and the apoptosis marker cleaved caspase3 (C-casp3) in MDA-MB-231 tumors after treatments. Scale bars = 10 μm. (**D**) EGFR protein expression in whole tumor lysates. Samples were collected from five different biological replicates. (**E**) Evaluation of acute liver and kidney parameters after a single dose of NTZ-NPs (10 mg/kg) or controls administered via the tail vein. Blood samples were collected and tested at 12 h post-injection. The heatmap represents the mean of three biological replicates. (**F**) DiR fluorescence signal in mice bearing transplanted U251 glioma after treatment with DiR dye loaded NTZ-NP or Ang/NTZ-NP. Angiopep-2 (Ang), as a tumor-targeting peptide and BBB-penetrating ligand, was conjugated to PEG-DSPE and mixed with NTZ-PEG-DSPE at a ratio of 1:4. (**G**) Statistics for 3 biological replicates of the DiR fluorescence signal in the mouse brain region.

To further confirm the potential for TPD-NP development as a therapeutic agent, we evaluated acute effects in the liver and kidney after TPD-NP treatment by measuring the following blood biochemical parameters. These data showed only slight changes (Figure 4E), indicating acceptable systemic response and biocompatibility. To further address the potential of TPD-NP for synergistic drug therapy, we loaded the near-infrared dye DiR into the TPD-NP to evaluate its drug loading and delivery capacity. Dye loading also allowed us to assess the performance of TPD-NP facing challenges in biological barrier penetration and tumor targeting. Moreover, to provide TPD-NPs with a passport to the brain through the blood‒brain barrier (BBB) and target glioma, we employed the angiopep-2 (Ang)-decorated PEG-DSPE arm with the NTZ-PEG-DSPE arm when performing self-assembly (Figure 4F) as Ang binds to the LRP-1 receptors expressed by endothelial cells of the BBB and glioma cells. After intravenous injection of the enhanced TPD-NPs, an increased DiR fluorescence signal was detected in U251 human glioma-bearing nude mice (Figure 4, F and G), suggesting good cargo loading and delivery capacities, and the potentials to readily equip TPD-NPs with biological barrier penetration ability and tissue targeting ability towards combination therapy.

## DISCUSSION

Extracellular TPD is now in the limelight of protein elimination-based therapy. However, the design and development strategy of current TPD tools requires laborious case-by-case design and *de novo* synthesis that need specific motifs to facilitate POIs cellular internalization and intracellular degradation. Our NP-based TDP strategy releasing from this constraint, may revolutionize the current TPD tools development landscape for clinical applications.

Multiple future directions may arise from this study. The frame of TPD-NPs is highly flexible. The biodegradable polymer-lipid hybrid NP used in this study is a general formulation for NPs. The PLGA core is not inherently essential and can be replaced with biodegradable polymeric materials that allow sufficient cellular uptake. Other forms of NPs, including polymersomes, micelles, DNA nanostructures, nanogels and inorganic NPs together with the biophysical properties (i.e., shape, size, surface potential and water solubility) of the TPD-NPs can be optimized for various applications.

TPD-NPs are highly feasible to synthesis through “mix-and-match”, and the POI hijack moiety and POI cellular location also have no strict limitations. Thus, TPD-NPs offer vast potentials for modification to suit a wide range of applications. Compared to new drug screening or synthetic modification of an existing drug as well as other existing TPD tools, the flexible superiority of the “nanoplatform” may greatly reduce R&D costs and time.

Another direction is “at-site” functional dose optimization towards pharmaceutical transformation. TPD-NP is superior *in vivo* because all the advantages of nanomedicines in terms of pharmacokinetics and pharmacodynamics. And plausible ways can further improve the potency (e.g adjusting the physicochemical properties of nanoparticles, lowering the steric hindrance, reducing antibody-PEG ratio).

Importantly, by employing the unique autolysosome degradation pathway and flexible sizing potential of TPD-NPs, TPD-NPs can be designed to match the size of large subset of misfolded/aggregated POIs ^29^ which may circumvent the “ant against elephant” problem when hijacking large aggregated proteins, such as β-amyloid oligomer or fiber, in neurodegenerative diseases. On the contrary, the lysosomal pathway and proteasome degradation pathway used by PROTACs has inherent size and substrate-conformation limitations and oligomeric proteins cannot be degraded and may even choke the pores of proteasomal subunits ^30^.

Moreover, our TPD-NP archetype, by exploiting basic knowledge in nanodelivery, enables facile cell-type-specific targeting, drug loading, and biological barrier penetration. Nanotechnology is changing the landscape of pharmaceutical industries and challenging traditional pharmaceutical science by offering such abilities. Our NP-mediated TPD mechanism may find broad applications with further rational design and may have exciting translational potential.

Fundamentally, the novelty of this study goes beyond TPD. As ligand-mediated tissue targeting strategies for nanoparticle-based delivery involve surface ligand engulfment, it is possible that the targeting efficacy may be reduced by a continuously high frequency of administration. This may be useful knowledge as principle to consider in the design of active nanomedicine protocols. Thus, it sheds new insights on receptor mediated drug therapies.

## ACKNOWLEDGMENTS

Schemes in figures were created with Biorender.com. National Natural Science Foundation of China (NSFC 52073079, 32071388) National Key Technologies R&D Program of China (2018YFA0209800)

Program of Technology Innovation Team in Colleges and Universities of Henan Province (21IRTSTHN028)

NHMRC Investigator Grant

## AUTHOR CONTRIBUTIONS

Conceptualization: MZ, YL, BS Methodology: YL, MZ, BS

Investigation: YL, RL, SW, HY, LF, MF, JD Visualization: YL, RL, XX, YZ

Funding acquisition: MZ, BS Project administration: MZ, BS Supervision: MZ, BS

Writing – original draft: YL

Writing – review & editing: MZ, BS, KWL

## DATA AND MATERIALS AVAILABILITY

All data are available in the main text or the supplementary materials.

## Supplementary Materials

Materials and Methods

Supplementary Data

