## Supplementary material for "Targeted Protein Degradation via Nanoparticles": Methods and Supplementary Data

### METHODS DETAILS

#### Synthesis of antibody-PEG-DSPE conjugates

We nonspecifically labeled lysine residues on antibodies with NHS-PEG-DSPE, generating the antibody-PEG-DSPE chain. We first replaced the original antibody buffer with PBS. For example, cetuximab solution was concentrated with a 3 kDa spin filter (Millipore) at 4000×g for 2 min and subsequently diluted with PBS, and the protein concentration was measured through the IgG mode of a NanoDrop One C and verified by a BCA protein concentration assay kit. Then, the antibody and NHS-PEG-DSPE conjugation reaction was performed in a $N_2$-containing bottle in an ice-water mixture environment, where an equal molarity and volume of NHS-PEG-DSPE in PBS buffer after sonication at 40 kHz for 3 min was added to the antibody solution with stirring at 800 rpm (revolutions per minute). The stirring reaction was then allowed to incubate for 24 h at 4 °C on a 20 rpm rotator. The reaction mixture was then filtered 3 times with a 50 kDa centrifugal filter (Millipore) and resuspended in 100 μL of 1×PBS, and the protein concentration was determined.

#### Synthesis of Peptide-PEG-DSPE

We conjugated the free $NH_2$ residues on peptides with NHS-PEG-DSPE at a certain molarity. The peptide powder and equivalent (molar ratio) NHS-PEG-DSPE powder were dissolved in 2 mL of DMF under the protection of nitrogen, and then 1.5 equivalents of trimethylamine were added. After 12 h of stirring at 800 rpm in a 37 °C water bath, the reaction solution was dialyzed through 2 kDa for 12 h to remove the unconjugated materials, and then the product was lyophilized.

#### Preparation of NPs

For the antibody/peptide-PEG-DSPE-(PLGA) shell-core NP formulation, we employed a self-assembly method to prepare polymer–lipid hybrid NPs as previously described (Islam et al., 2018). Briefly, PLGA (15 kDa) dissolved in DMF was dropped dropwise (2 μL per drop, 5 sec per drop) into the 600 rpm stirring NTZ-PEG-DSPE solution (molar ratio PLGA: NTZ-PEG-DSPE=1:1) and stirred for 30 min at room temperature. Then, the solution was concentrated by a 2 mL 3 kDa ultrafiltration tube at 12000 g 5 times for 4 min each time to remove the unconjugated materials. Then, the protein concentration was measured by a NanoDrop One C (Thermo Fisher) in IgG mode and verified by a BCA protein assay kit. Exosomes was extracted by ultra-centrifuge method from mouse dendritic cell line DC2.4 cell line according to general methods and calculated by protein concentration. The liposomes used in this study was composed of HSPC, CHO through a classic filming-rehydration method, where the elements were weighed accordingly to the weight ratio of HSPC: CHO = 56: 39, and then dissolved in chloroform, and condense the liposome to a membrane by evaporation. After complete evaporation of chloroform, the Antibody/peptide–PEG-DSPE arm in PBS buffer was added for resuspension, followed by sonication at 100 W at room temperature for 3 min. The resulted solution was then filtered through 800 nm, 400 nm and 200 nm filter membranes to obtain the even size. The weight ratio of HSPC: CHO: PEG-DSPE = 56: 39: 2.5 was modified refer to the liposomal drug Doxil.

**Cell culture**

Cells were grown in cell culture dishes (Wuxi NEST Biotechnology, Ltd, China) under standard incubation conditions at 37 °C in the presence of 5% $CO_2$. The established cell lines MDA-MB-231, HeLa, U87 and U251 were maintained in DMEM high glucose 4.5 g/L medium, and HepG2 was maintained in DMEM low glucose 1 g/L medium supplemented with 10% fetal bovine serum (FBS) (Cytiva Lifesciences), 1% penicillin–streptomycin solution (Gibco), nonessential amino acids (Gibco) and Glamax (Gibco). When the confluency of the cells reached 90%, the cells were detached and passaged with 0.25% trypsin containing EDTA (Gibco). Transient plasmid transfection was carried out by Lipofectamine 2000 (Invitrogen) according to the manufacturer's instructions. Cells were checked for mycoplasma contamination-free status before *in vitro* cell experiments and *in vivo* xenograft tumor model preparation. Cell viability was measured by the CCK-8 method.

**Physicochemical characterization of the NPs**

The synthesized NPs were characterized by a dynamic light scattering system (DLS, Zetasizer Nano-ZS, Malvern Instruments) and transmission electron microscopy (TEM, JEM-2010HT Japan). The TEM samples were prepared on copper grids and dried. Then, the sample-containing grids were stained with 1% uranyl acetate for 5 min at room temperature, washed with deionized water 3 times and imaged by TEM at 120 kV. Successful antibody-NP conjugation was verified by Coomassie blue staining in a 4-20% gradient native gel after electrophoresis.

**Flow cytometry assay**

For the surface protein expression assay, cells were seeded into a 12-well plate with $1\times10^5$ cells per well, and 500 μL medium was supplied. Cells transfected with the C-terminal fused GFP tag were lifted with EDTA (Gibco, 15040066) and washed 3 times with PBS (containing 0.5% BSA). Then, flow cytometry was performed on a CytoFLEX (Beckman) flow cytometer, and FlowJo software was used to analyze the data. Gating was based on single cells and live cells.

**Immunostaining and microscopy**

Cells used for confocal imaging were seeded onto a confocal imaging coverslip in a 12-well plate ($1\times10^5$ cells per well). For surface protein expression, cells were washed 3 times with PBS (containing 0.5% BSA), fixed with 4% paraformaldehyde for 15 min in the dark, incubated with 1% BSA PBS solution for 1 h and washed with PBS. Then, cells were incubated with primary antibody overnight at 4 °C, followed by 3 washes with PBS (containing 0.5% BSA). Then, secondary antibodies were incubated for 1 h at room temperature in the dark. After incubation, the cells were washed with PBS, stained with DAPI and imaged with a Zeiss 880 confocal microscope. For live cell imaging, cells were cultured in a cover glass bottom dish and visualized by a Zeiss 880 confocal microscope with a cell culture system to ensure that the culture was at 37 °C in the presence of 5% $CO_2$ and in a humid environment. For tissue immunofluorescence staining, fresh tissue was collected, fixed with 4% paraformaldehyde overnight, and embedded in paraffin. Tissue resection was 5 μm and was permeabilized by incubation in 0.2% Triton X-100-PBS for 10 min followed by blocking with 2% serum in PBS for 1 h at room temperature. Next, the samples were incubated in primary antibody at 4 °C overnight, followed by PBST washing and secondary antibody incubation for 1 h at room temperature. Then, slices were counterstained with 500 nM 4,6-diamidino-2-phenylindole (DAPI) and mounted with Prolong Gold antifade mounting medium (Invitrogen). The cells were imaged with a Zeiss 880 confocal microscope.

**Animal model**

All the animals used in this experiment were specific pathogen free and provided by SiBeiFu Laboratory Animal Technology Co. Ltd. Mice were fed for at least 1 week to acclimatize to the food and environment, and automatically controlled animal facilities with a temperature of 23±2 °C and humidity of 50 ± 20% under a 12-h light/dark cycle were provided. Mice were provided free access to sterile pellet food and water. All animal experimental protocols were approved by the Henan University Laboratory Animal Centre and the Animal Care and Use Committee of Henan University (Approval ID: HUSOM-2022-195) and performed under standard guidelines.

**TPD-NP performance evaluation in the MDA-MB-231 xenograft tumor model**

To prepare the MDA-MB-231 xenograft tumor mouse model, $4\times10^6$ MDA-MB-231 cells in 200 μL Matrigel (BD Biosciences) containing culture medium (1:1) were implanted subcutaneously into 6- to 8-week-old female athymic nude mice. Mice were monitored for tumor volume every day, and when the majority tumor size of mice reached 100±10 $mm^3$, mice with tumor size 100 ± 10 $mm^3$ were randomly grouped (10 mice per group). Then, 15 mg/kg (antibody/body weight) antibody or NP conjugates in 100 μL PBS were injected into each group every second day, and the tumor volume was recorded every day. Ellipsoid tumor volume was calculated by the formula: Volume = $\pi/6\times\text{Width}^2\times\text{Length}$.(Tomayko and Reynolds, 1989) All treatments were performed according to the antibody amount, and the IgG protein concentration was verified by a NanoDrop one C. All NPs used were prepared and characterized before injection. Mice were sacrificed at the indicated time points, and the tumor sizes were recorded and further prepared for Western blots or immunostaining.

**Blood assay and bioimaging**

Ten milligrams $kg^{-1}$ of antibody or NP conjugate was intraperitoneally injected into female 6–8-week-old BALB/c mice (3 mice per group). The grouping of mice was random. At the indicated time points, blood was sampled from the eye socket using anticoagulant capillary tubes, and serum was separated after centrifugation at 700 g at 4 °C for 15 min. Then, the blood chemistry parameters for major liver and kidney metabolism were analyzed using a kit from Wuhan Servicebio Technology Co., Ltd. To evaluate the *in vivo* tumor targeting and blood–brain barrier penetrating ability of TPD-NPs, Ang/PEG-DSPE was synthesized as previously described by reacting angiopep-2 (Ang) $NH_2$ residues with NHS-PEG-DSPE and assembled in a mix-and-match manner, in which 20% Ang-PEG-DSPE was mixed with NTZ-PEG-DSPE and precipitated onto the PLGA core. A GBM-bearing nude mouse model was prepared in which fully anesthetized mice underwent the brain operation. U251 human glioma cells were injected into the brain and sealed with tissue glue, and the mice were kept in a warm environment following the standard protocol. The TPD-NPs were injected intravenously into GBM-bearing nude mice and monitored by using the Lumina IVIS III Imaging System (DiR dye wavelength excitation=640 nm; emission=690 nm).

**Western blot assay**

Western blot assays were performed according to a standard protocol to detect protein expression. The protein bands were visualized by Amersham Imager 680 (General Electric Company). For sample preparation, cells were seeded in a 12-well plate ($1\times10^5$ cells per well), and treatments with TPD-NP at different doses were diluted to the medium before being added to the culture medium to a final volume of 500 μL. The dose usage for the control groups referred to the conjugated antibody/peptide/small molecular POI binder. Treated cells were harvested with RIPA lysis buffer containing protease/phosphorylase inhibitor cocktail at the indicated time points on ice by a cellular scraper after 3 washes with PBS. The collected cell lysates were freeze-thawed 3 times to mechanically break the cellular structure and then centrifuged for 10 min (15,000 rpm, 4 °C). After centrifugation, the supernatant was collected, and the protein concentration of the supernatant was determined using a BCA protein analysis kit.

**Statistical analysis**

All graphs and statistical analyses were carried out using GraphPad Prism 8 software. Two-sided t tests and one-way analysis of variance (ANOVA) were used. All experiments were performed in triplicate unless otherwise stated. Error bars indicate the standard error of the mean (SEM) unless otherwise noted as the standard deviation (SD). The $P$ value < 0.05 was considered statistically significant, whereby all significant values shown in various figures are indicated as follows: *$P < 0.05$, **$P < 0.01$ and ***$P < 0.001$.

### Supplemental figures

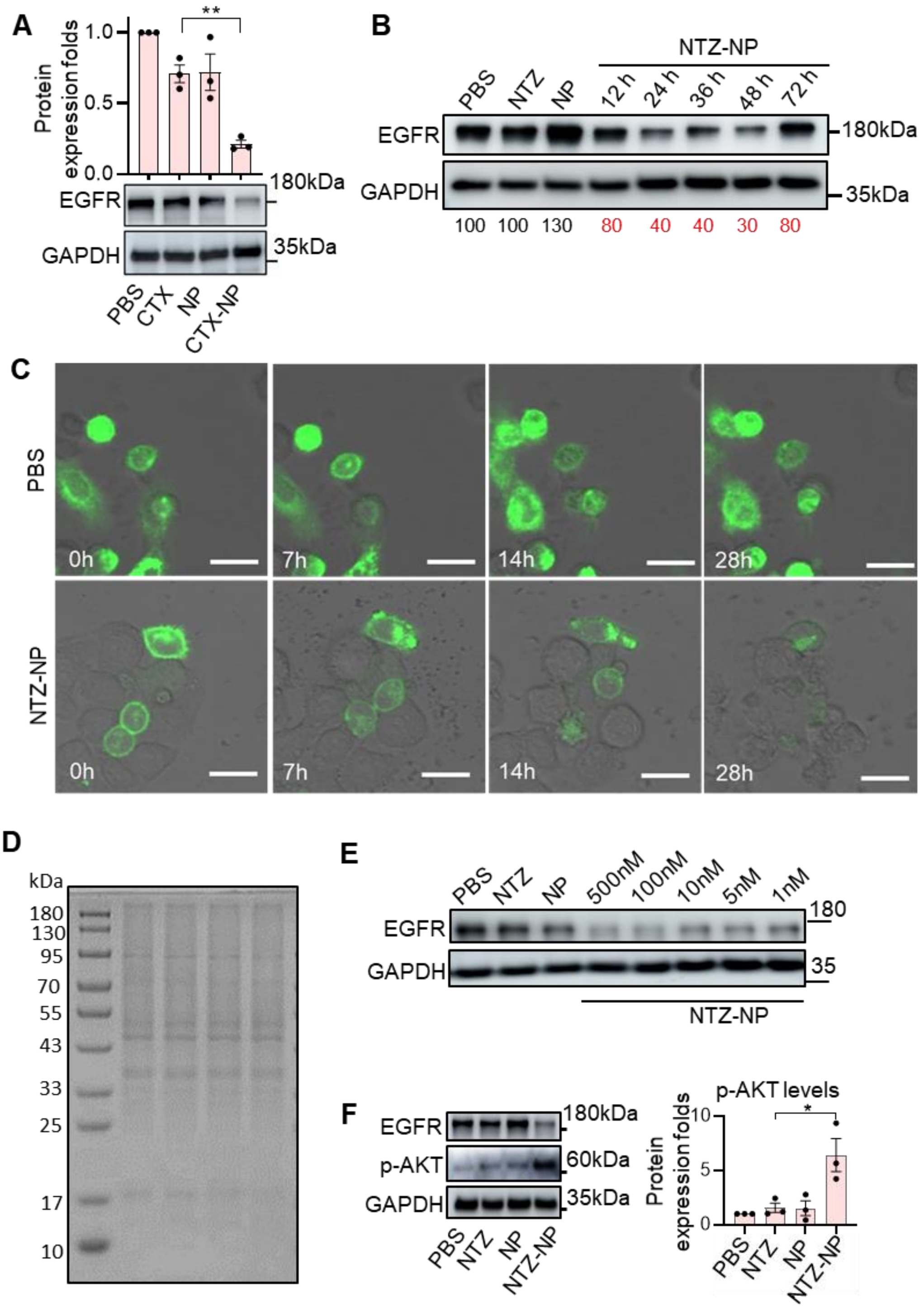


**Figure S1. Dose challenging, reversible POI degradation manner, degradation time series, global protein changes and key down stream of EGFR response for NTZ-NP treatments. Related to Figure 1**

(A) EGFR protein expression and statistics of Western blot bands correlated with its GAPDH, samples collected from MDA-MB-231 cells treated with 500nM CTX-NP for 24h. (CTX, cetuximab, clinical approved EGFR monoantibody).

(B) NTZ mediates a reversible EGFR degradation in a time dependent manner, indicating the TPD do not disturb protein recycling. MDA-MB-231 cells were treated with 500 nM NTZ-NP for 24h, then the culture medium was removed and washed, then fresh medium was supplied, sample was collected at indicated time points after treatment. These data provided a basic knowledge for the nanomedicine dosing strategy design, that the ligand can mediate surface receptors degradation.

(C) Degradation of cell surface EGFR in MDA-MB-231 cells as determined in live cells with fixed visual field after treatment with 500 nM equivalent antibody dose of NTZ-NP. MDA-MB-231 cells were transfected with the EGFR-EGFP vector, where the EGFP tag was fused to the intracellular domain (C-terminal before the stop codon) of EGFR. Scale bar = 25 µm. Images also were made into videos.

(D) Native gel with Coomassie staining of the NTZ-NP treatments (24h, 500nM, MDA-MB-231 cells).

(E) NTZ linked TPD-NP mediates EGFR degradation in a dose dependent manner. (24h, dose was referred to antibody dose, MDA-MB-231 cells). Grayscale statistics for EGFR bands correlated with GAPDH was noted. Protein degradation can be achieved at least 1 nM, which performance is competitive to minimal dose 10 nM LYTAC (*Nature* 2020).

(F) Upregulation of active phosphorylated-AKT (p-AKT) as the complementary HER3/IGF1 pathway downstream in response to EGFR degradation was reported and validated. Statistics were from three biological replicates. $P<0.05$, mean ± s.e.m.

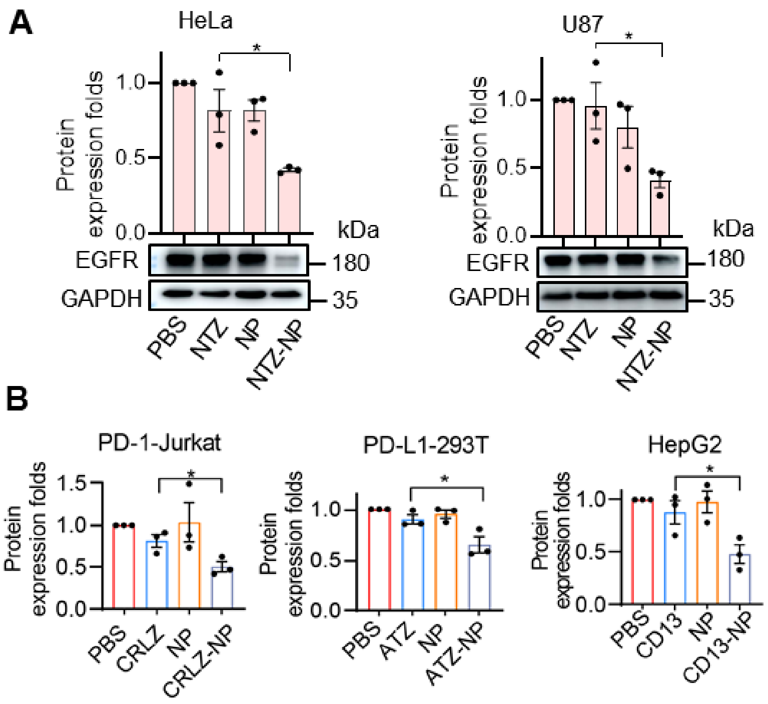


**Figure S2. POI degradation after TPD-NP treatments in different cell types. Related to Figure 2**

(A) EGFR protein levels after treatment with 500 nM NTZ or NTZ-NP conjugates for 24 h in HeLa cells and U87 cells.

(B) Statistics for targeted protein bands correlated with its GAPDH. Cells were treated with different NP conjugates for 24 h. Representative Western blot data was presented in Figure 2.

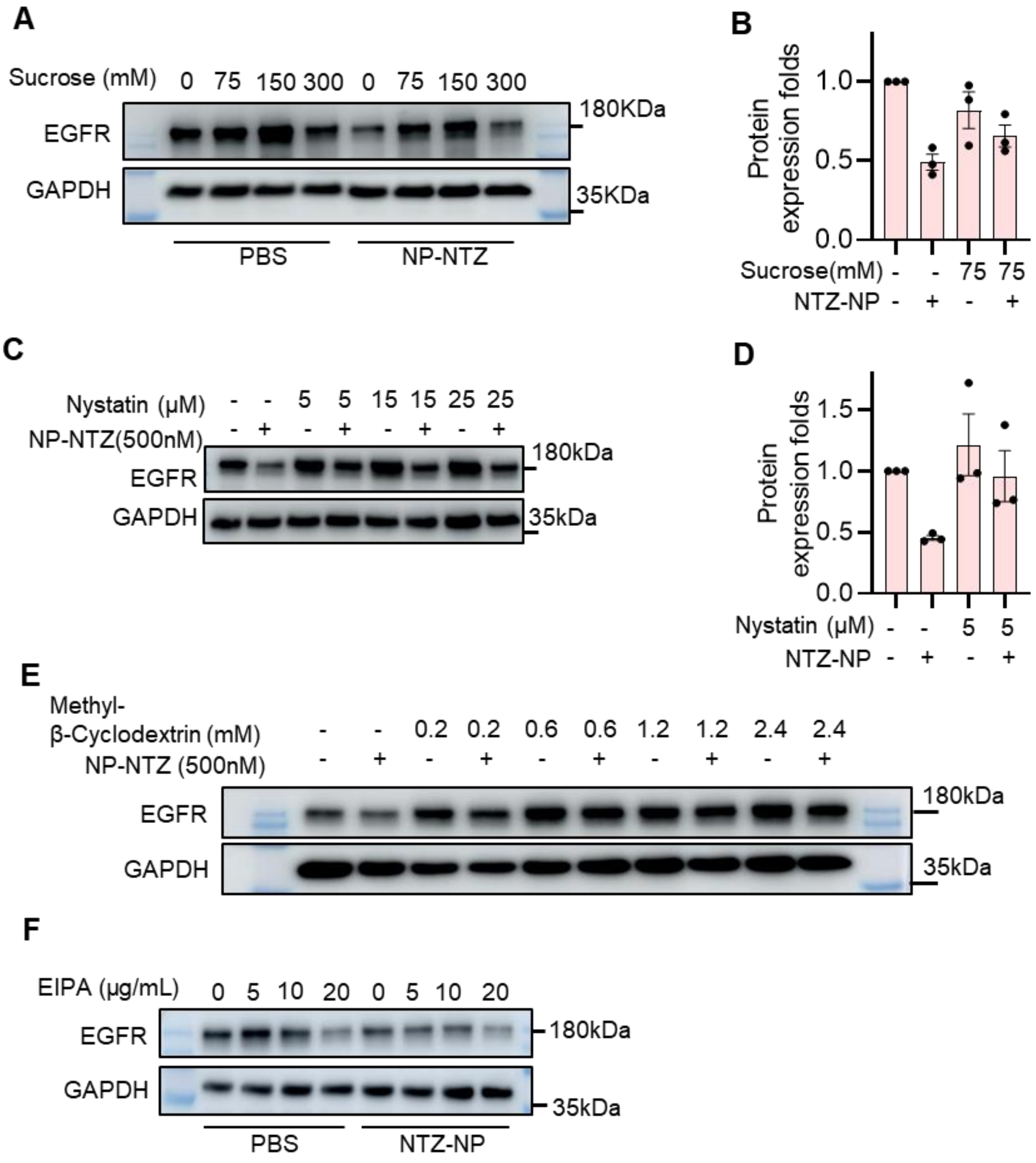


**Figure S3. Inhibitors dose series and statistics for the endocytosis pathway. Related to Figure 3.**

(A) EGFR expression in MDA-MB-231 after treatment with different dose of sucrose (clathrin-mediated endocytosis inhibitor) for 24h.

(B) Statistics for the EGFR band grayscale correlated with the GAPDH after treatment with Sucrose and NTZ-NP for 24 h. n=3. mean±s.e.m.

(C) EGFR expression in MDA-MB-231 after treatment with 500 nM NTZ-NP and different dose of nystatin (caveolin-mediated endocytosis inhibitor) for 24h.

(D) Statistics for the EGFR band grayscale correlated with the GAPDH after treatment with Nystatin and NTZ-NP for 24 h. n=3. mean±s.e.m.

(E) EGFR expression in MDA-MB-231 after treatment with 500 nM NTZ-NP and different dose of Methyl-β-cyclodextrin, (MβCD, caveolin-mediated endocytosis inhibitor).

(**F**) Expression of EGFR protein in MDA-MB-231 cells treated with 500 nM NTZ-NP and different concentration of EIPA (macropinocytosis inhibitor) for 24h.

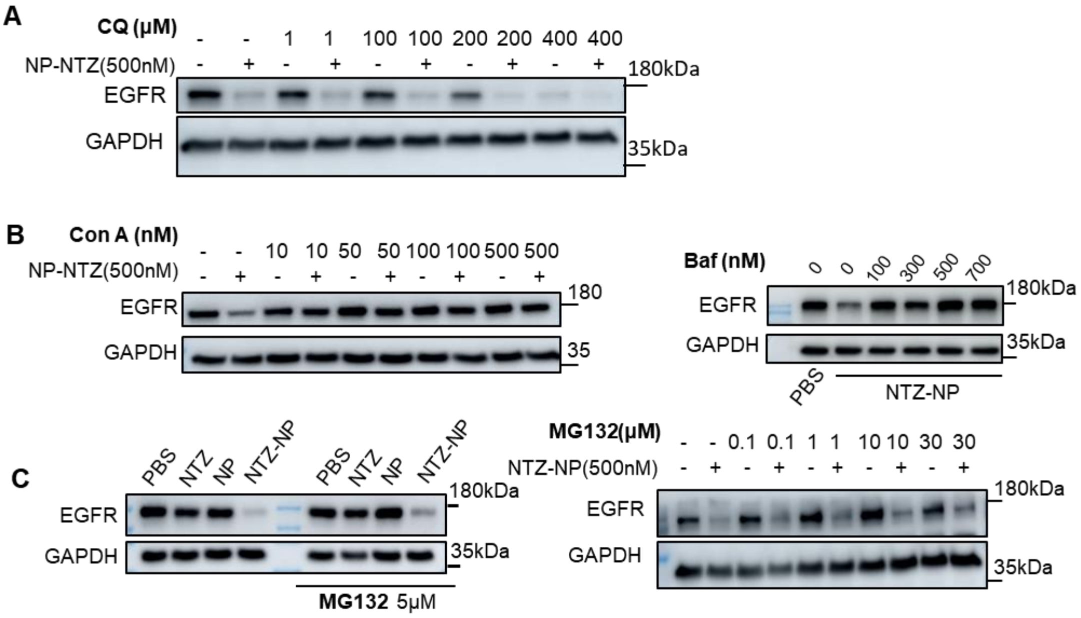


**Figure S4. Inhibitors dose series for the protein degradation pathway. Related to Figure 4**

(A) Expression of EGFR protein in MDA-MB-231 cells treated with different doses of chloroquine (CQ, lysosome inhibitor) for 24h.

(B) Expression of EGFR protein in MDA-MB-231 cells treated with different doses of concanamycin A ($H^+$-V-ATPase inhibitor, inhibiting the acidification of the autolysosome) for 24h. Treatment with different doses of baflomycin A1 (Baf, $H^+$-V-ATPase inhibitor) was also shown, right panel.

(C) Expression of EGFR protein in MDA-MB-231 cells treated with 500 nM NTZ-NP and different doses of MG-132 (proteasome inhibitor) for 24h. EGFR band grayscale correlated with the GAPDH was noted. MG-132 over 30 μM showed a decrease in cell viability, 30 μM or less of MG-132 fails to inhibit the EGFR protein degradation.

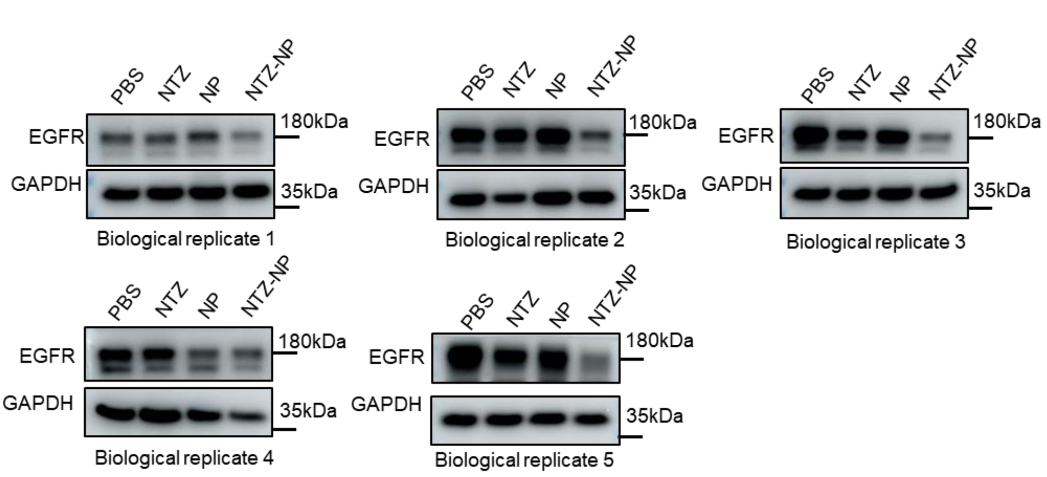


**Figure S5. Western blots used for EGFR expression statistics in MDA-MB-231 subcutaneous**

**tumor transplantation model. Related to Figure 4**

Samples in each biological replicate group are collected from whole tumor lysate from total 20 mice with different treatment.

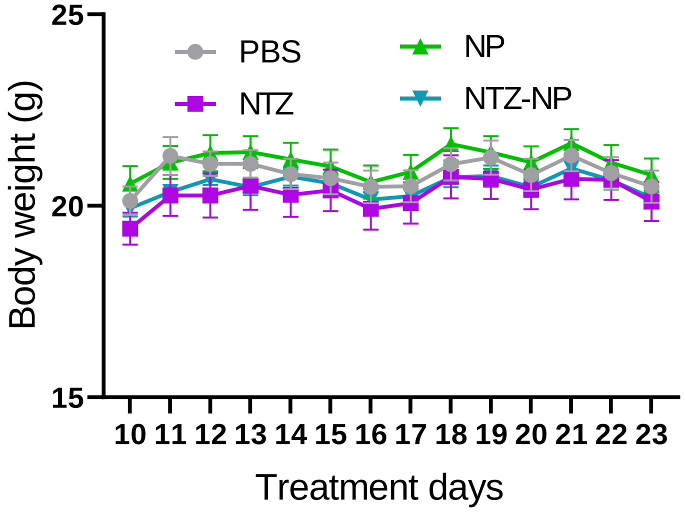


**Figure S6. Body weight changes during MDA-MB-231 subcutaneous tumor transplantation model. Related to Figure 4**
